# Broadscale reconnaissance of coral reefs from citizen science and deep learning

**DOI:** 10.1101/2024.11.27.625580

**Authors:** Christopher L. Lawson, Kathryn M. Chartrand, Chris M. Roelfsema, Aruna Kolluru, Peter J. Mumby

## Abstract

Coral reef managers require various forms of data. While monitoring is typically the preserve of scientists, larger scale reconnaissance data that can be used to inform spatial decisions does not usually require such precise measurement. There is an increasing need to collect such broadscale, up-to-date environmental data at massive scale to prioritise limited conservation resources in the face of global disturbances. Citizen science combined with novel technology presents an opportunity to achieve data collection at the required scale, but the accuracy and feasibility of new tools must be assessed. Here we show that a citizen science program that collects seascape images and analyses them using a combination of deep learning and online citizen scientists can produce accurate benthic cover estimates of key coral groups. The deep learning and citizen scientist analysis methods had different but complementary strengths depending on coral category. When the best performing analysis method was used for each category in all images, mean estimates from 8086 images of percent benthic cover of branching *Acropora*, plating *Acropora*, and massive-form coral were ∼99% accurate compared to expert assessment of the same images, and >95% accurate at all coral cover ranges tested. The effort to achieve 95% accuracy at a site – our ecologically relevant target based on the accuracy of other tools – was attainable based on citizen scientist involvement in pilot years of the program, with 18-80 images needed depending on coral type and reef state. Power analyses showed that sampling up to 114 images per site was needed to detect a 10% absolute difference in coral cover per category (power = 0.8), accounting for natural heterogeneity. However, the benthic cover of ‘all other coral groups’ as a single category could only be estimated with 95% accuracy at 60% of survey sites and for images with 10-30% coral cover. Disaggregating this ‘other coral’ group into more distinct coral categories may improve accuracy. Overall, citizen science can provide an accuracy that is acceptable for many end-users for select coral morphologies. Such a combination of emerging technology and citizen science presents an attainable tool for collecting inexpensive, widespread reconnaissance data of coral reefs that can complement higher resolution survey programs or be an accessible tool for resource-poor locations.

## Introduction

Ecosystem management needs various forms of data (Grêt-Regamey et al., 2017; Lindenmayer et al., 2008). Long-term monitoring of coral reefs is often conducted by government and research programs focused on accurate estimates of coral abundance at high taxonomic resolution (Edmunds, 2024; Reverter et al., 2022). However, there is also a need for coarser, rapid reconnaissance over large areas (Edmunds & Bruno, 1996; Mumby et al., 2021). Such up-to-date broadscale reconnaissance will inform where to prioritise limited conservation resources in the face of unprecedented global disturbances (Reverter et al., 2022; Swinfield et al., 2024).

One method to achieve broadscale reconnaissance is citizen science, whereby effort is crowdsourced from distributed participants. Citizen science has contributed data on coral reefs for decades. In the 1980’s, Raleigh International conducted dedicated project-based expeditions and marine surveys by trained citizen scientists (Beames, 2004). In the 1990s, Coral Cay Conservation trained citizen scientists to collect data in support of establishing coral reef management plans in Belize (Mumby et al., 1995). More recently, Reef Check engages trained citizen scientists to capture percent cover of 10 benthic cover categories using point intercept transects; it collects data that are ∼93% accurate and aims to support science and management decisions (Done et al., 2017; Hodgson, 1999). Established in 2007, Reef Life Survey uses selectively chosen and trained citizen scientists to collect high-quality on Scuba, supporting global science and conservation efforts (Edgar & Stuart-Smith, 2014).

There are also government-run citizen science programs such as Reef Health Impact Surveys and Eye on the Reef, operated by the Great Barrier Reef Marine Park Authority in Australia (Beeden et al., 2014). Reef Health Impact Surveys provide ‘advanced in-water training’ to citizen scientists to collect data in a structured program. The Eye on the Reef mobile application is simpler and relies on opportunistic sampling that enables observational data collection by anyone on the Great Barrier Reef.

The CoralWatch citizen science program was established in 2002, creating a simple tool to assess the presence of coral bleaching by comparing *in-situ* coral colour with a calibrated coral health chart. CoralWatch differs from many previous programs because it does not require substantial training and enables anybody to collect data, resulting in a large, opportunistically collected database (Marshall et al., 2012). CoralWatch currently comprises 17% of all publicly accessible bleaching data globally through its online data portal (unpublished data, C. Roelfsema).

Some of the limitations for citizen scientists to participate in accurate data collection may be removed by using technology such as deep learning (McClure et al., 2020). Deep learning, a subdomain of artificial intelligence, is a computational approach in which systems learn patterns from data, rather than following explicit instructions, enabling them to solve tasks based on examples rather than pre-defined solutions (Mitchell, 1997). Deep learning has dramatically increased the efficiency of environmental image analysis (e.g. González-Rivero et al., 2020). However, current deep learning tools for coral reefs mostly rely on consistent, high quality photographs of quadrats (Courtney et al., 2022; González-Rivero et al., 2020; Schürholz & Chennu, 2023). While such photographs could be taken by citizen scientists, it requires dedicated Scuba logistics, which best suits the capacity of professional scientists engaged in monitoring reef state. Opening image collection to citizen scientists without training, specialist equipment, and with flexible logistics including snorkelling, would vastly expand the scope of data collection.

The Great Reef Census is a citizen science project that started on the Great Barrier Reef, Australia. The Great Reef Census utilises two types of citizen scientist: those who collect underwater images in the field and those ‘virtual volunteers’ who help analyse the resulting images online. The latter group are based all over the world and do not need access to the reef: many do not have access due to distance, resources or physical limitations. For in-water field surveys, citizen scientist tourists and reef industry workers capture images without specialised equipment or formal training. The only training required is reading a simple 2-page methods protocol. These images are then analysed using deep learning and by online citizen scientists to estimate benthic cover. A key question is if using deep learning reduces the barrier to entry for non-experts to participate in basic image analysis. Deep learning is generally faster at recognising shapes and is rapidly improving, but human vision may still outperform when complexities are introduced such as texture, shadows or poor water visibility (Rubbens et al., 2023).

There is a need to assess if citizen science-based seascape photo analysis can provide valid data to inform management, restoration or science. If image collection can be achieved by nearly anyone and analysis can be distributed to deep learning (artificial intelligence; hereafter ‘AI’) and citizen scientists globally, this would enable a vast expansion of the scope of data collection relative to traditional tools. However, achieving massive scaling of data collection requires a trade-off in precision, accuracy and taxonomic resolution. Because scale and accessibility for non-experts is limited by the complexity of species-level identification, here we do not identify specific taxonomies, which are constantly under revision and even beyond the skillset of many scientists (Ramírez-Portilla et al., 2022). Yet, measuring cover of select coral morphologies can still inform many management actions, such as pest control and marine park planning, and morphological information by genus is important for key ecosystem functions like bioconstruction of reefs (Wolfe et al., 2020). Here, we focus on the capacity of citizen science to estimate cover of key coral morphologies that commonly dominate on the Great Barrier Reef: branching *Acropora*, plating *Acropora* and massive-form corals such as *Porites* or *Platygyra* (Veron, 2000). Branching and plating *Acropora* are fast-growing coral that are important for reef recovery following disturbance, but are vulnerable to threats like crown-of-thorns starfish and cyclones, while massive corals are slower growing yet more resistant to threats and exhibit longevity that is important for sustaining reef accretion and persistence (Loya et al., 2001; Ortiz et al., 2021; Pratchett et al., 2020; Wolfe et al., 2020). Protecting populations of these coral groups can give outsized ecological benefit (Ortiz et al., 2021).

Our overall aim is to assess if seascape images of the reef collected by citizen scientists can provide sufficiently reliable information for reef management. To achieve this aim, our first objective is to assess if AI-alone or AI-supported citizen scientist analysis can accurately quantify the cover of three coral groups in seascape images collected by citizen scientists. Next, given the variability in accuracy among images, we ask how many images are needed to achieve a reasonable level of accuracy for a survey site, and how many online citizen scientists are needed to analyse each image. Finally, we run a series of power analyses to determine the number of images needed to also account for the natural heterogeneity of the reef.

## 1. Methods

### 1.1. Image collection and analysis

We analysed seascape images collected by citizen scientists using three methods: a semantic segmentation deep learning model (‘AI-alone’), an AI-assisted online citizen scientist analysis platform (‘AI+Citizen’), and ‘expert’ analysis which was used to assess the performance of the other two methods. We then explored the performance of the results in deriving accurate coral cover values with current resource capabilities (Figure 1).

**Figure 1.**
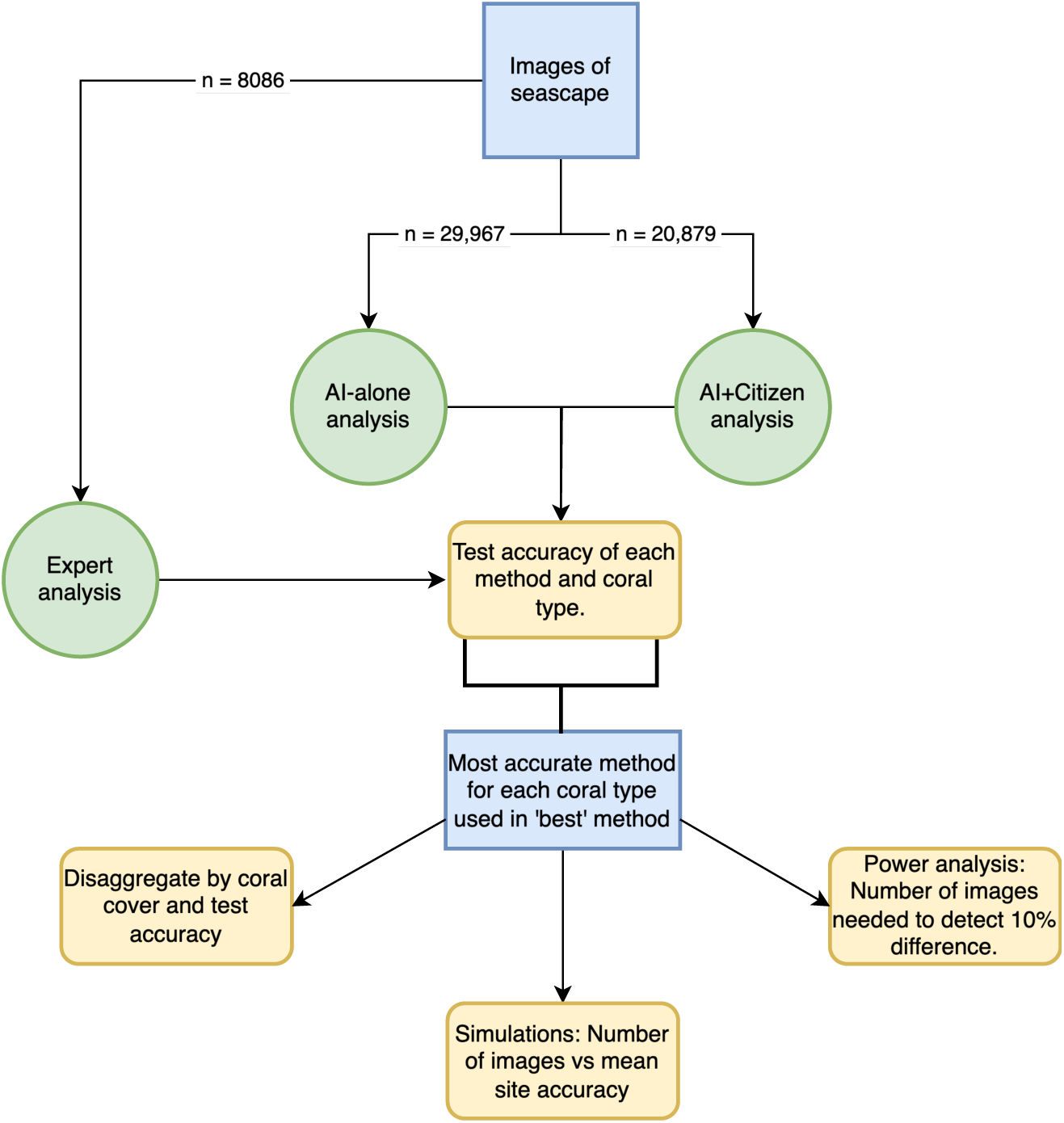
Summary flowchart of methods. Blue rectangles represent data, green circles represent methods of image analysis, and yellow rounded boxes represent statistical analyses of performance. Figure made with draw.io.

#### 1.1.1. In-water survey methodology

Images (n = 29,967) were collected as part of the Great Reef Census from September 2022 – February 2023 at 1512 sites distributed over 211 of the ∼3000 reefs across the Great Barrier Reef, Australia. The Great Reef Census follows a simple survey methodology to collect seascape images (See Figure 3 for an example). Volunteer citizen scientists (∼70) were tasked with capturing random images of reef slopes or bommies at depths between 3 m and 20 m. While participants could survey any reef, a priority map of reefs was provided to guide the most ‘valuable’ reefs to survey based on relevance to government managers, scientists or ecological importance, for example as a key source of larval dispersal (Mumby et al., 2021). Shallow reef tops (0-3 m) were excluded due to the difficulty of obtaining seascape images.

The survey protocol was designed to be easy, without the need for advanced training or scientific equipment. Images were collected on snorkel further from the substrate than standard photoquadrat surveys-i.e. 3-5m compared to 1m (Williams et al., 2019)-using basic handheld cameras such as GoPros (www.gopro.com). Images were captured parallel to the reef, with snorkelers duck-diving as required.

Participants were told to capture images every 10 fin kicks, worked in pairs, and aimed to photograph reef sections approximately 5 m × 5 m in each image, with a minimum of 20 images per person per survey. Participants were instructed to survey at least three sites of a reef, separated by a minimum distance of 200 m. Preferably each site was located on a different aspect of the reef, i.e. north, south, east/windward or west/leeward, assuming safe and feasible logistics. Images were uploaded to the Great Reef Census web-based platform (www.greatreefcensus.org) with corresponding time and GPS coordinates. GPS coordinates were given for each image if a towed GPS unit was used, otherwise GPS coordinates were noted at the beginning of each survey from the mother vessel, the tender vessel, or the camera’s internal GPS while it was above water.

#### 1.1.2. Expert validation data

To assess the accuracy of the AI-alone and AI+Citizen analyses, a subset of images were analysed with high accuracy using manual analysis by paid scientists skilled in coral identification and other benthic categories (hereafter referred to as ‘expert’ data).

To establish an efficient method of expert analysis, 615 images were first analysed by two methods: a ‘detailed’ method and a ‘visual’ method (Jokiel et al., 2015; Josephitis et al., 2012). The ‘detailed’ method used a custom-built software to draw polygons manually around individual coral colonies and assign a label corresponding to the coral categories of interest. The label options were branching *Acropora* (hereafter ‘Branching’), plating *Acropora* (hereafter ‘Plating’), massive-form coral (hereafter ‘Massive’), all other coral (hereafter ‘Other’)”, “reef substrate”, “water, sand, and shadow”, and “I don’t know” (Figure 2). The total area of each coral category’s polygons in each image were then calculated. Coral categories were presented as percent of total colonisable reef substrate, i.e. excluding sand/water/shadow. The ‘visual’ method used a different custom-built software that placed a 9-cell grid (3×3) over each image. Each grid square therefore comprised 11.1% of the total image. Experts visually assessed the proportion of each of the nine grid sections comprised of each coral category. The coral cover proportion of each grid square (0-100%) was multiplied by 11.1% and all grid square values summed to obtain the total cover of each coral type in each image. There was no significant difference in absolute coral cover between the ‘detailed’ and ‘visual’ methods (p = 0.6, mean difference =-1.5%, n = 615, Wilcoxon Signed-Rank Test).

**Figure 2.**
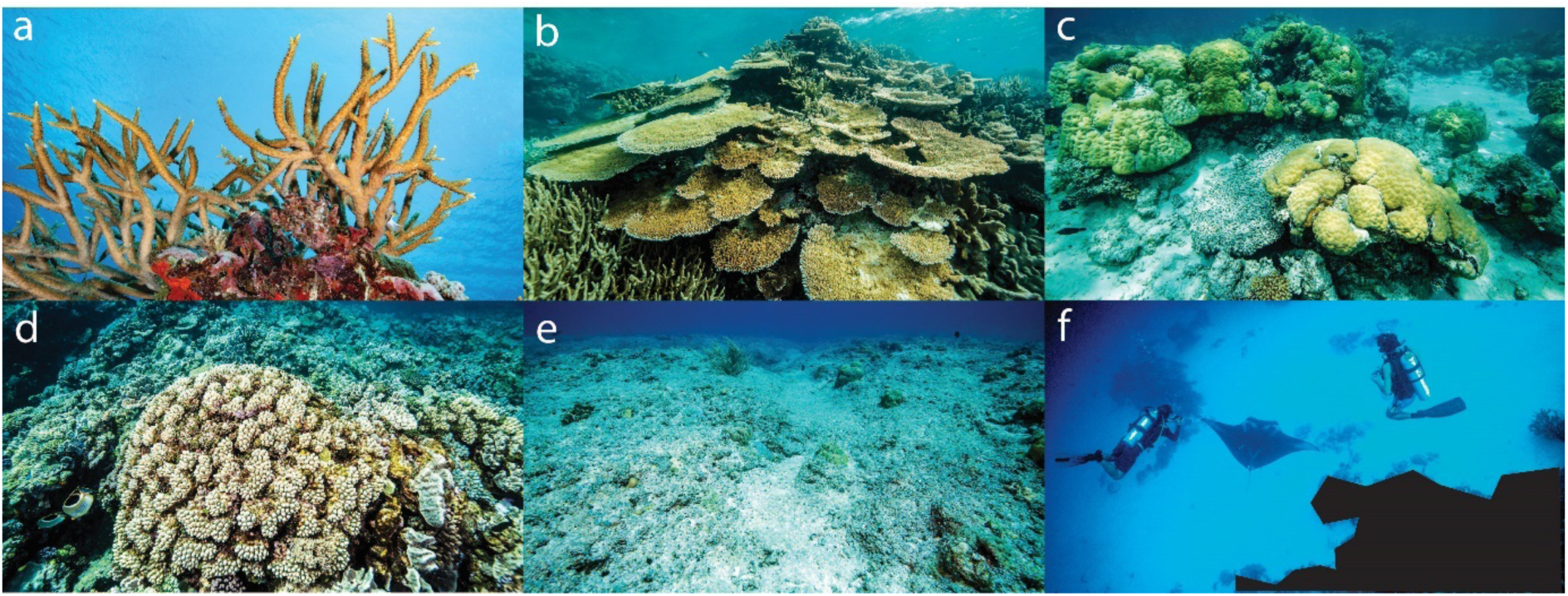
Category label options used for expert analysis, AI-alone analysis, and the AI+Citizens online analysis platform. A) “Branching Coral”-Branching coral of genus Acropora. B) “Plating Coral” - Plating/table coral of genus Acropora. C) “Massive Coral”. D) “Other Coral” - All other coral types. E) “Reef substrate” - any hard surface of the seascape suitable for coral growth. F) “Water, sand and shadow” - any region not included in the other categories, consisting of the background water column, bare sand, shadow or other objects that preclude substrate identification.

As a result, we used the faster ‘visual’ method to maximise the number of images analysed. Using the ‘visual’ method, 8086 images were analysed by three experts. Images were randomly assigned to experts; if the same image was analysed by multiple experts, the average values for each coral cover category were taken.

#### 1.1.3. Deep learning model development

A semantic segmentation model (Guo et al., 2018) was trained to identify coral morphology in citizen science imagery. SegFormer was used to develop the segmentation model (Xie et al., 2021). SegFormer uses a robust hierarchical transformer-based approach and its architecture allowed the model to capture fine-grained spatial features and contextual relationships within coral imagery. These characteristics are critical when analysing the variability in coral shapes, sizes, and colours, as well as the complex underwater environment with challenging lighting conditions and diverse backgrounds. The model was implemented in Python using PyTorch and trained on a Dell Technologies HPC GPU-Accelerated System, utilising a Dell EMC PowerEdge server cluster (Table 1).

**Table 1.** Parameter values used in training the semantic segmentation (AI) model.

| Parameter | Value | Comments |
| --- | --- | --- |
| Crop Size | 640 x 640 pixels | Provided a balance between computational efficiency and the preservation of crucial spatial features in the coral imagery. |
| Batch Size | 7 | Optimised memory usage on available hardware while ensuring stable gradient updates. |
| Learning Rate | 0.00006 | This relatively small learning rate was required for the fine-tuning process, enabling the model to gradually adjust to the intricacies of coral morphology without overshooting optimal parameter values. |
| Learning Rate Schedule | Constant, with no additional scheduling mechanisms | This approach was chosen after observing that the model's convergence was stable and that introducing a learning rate decay did not significantly improve performance during preliminary trials. |
| Maximum Epochs | 30 epochs | Determined through iterative experiments to ensure that the model had sufficient opportunities to learn while avoiding overfitting. |
| Early Stopping Patience Value | 20 epochs | Training would halt if no improvement in the validation mean Intersection over Union (IoU) score was observed over 20 consecutive epochs. |
| Early Stopping Mode | Maximum | Ensured that the best-performing model was retained based on IoU maximisation. |

To train the segmentation model, 7505 reefscape images collected by citizen scientists as part of Great Reef Census expeditions in 2020-2021 were annotated using the ‘detailed’ expert analysis method described earlier, using a custom-built software to delineate key coral morphologies digitally and assign labels to each polygon. The labels were the same predefined categories used in the expert analysis. The custom-built software converted these labelled polygons to JSON files used for segmentation model training (Table 1). The 7505 training images were divided in an 80:20 split: 6,004 images were used to train the model directly and 1,501 images were used for validation and evaluation to allow the model to learn effectively during training (Table 1).

After training, the model was used to generate segmentation masks of 29,967 images that weren’t involved in the training phase, classifying each pixel into one of the predefined categories. The model produced a total pixel count of each category that was divided by the known total pixel count of each image to determine percent cover of each category and used as the ‘AI-alone’ values for each image.

#### 1.1.4. AI+Citizen analysis platform

An online platform was created (www.greatreefcensus.org/analysis) where citizen scientists assign labels to polygons for each image to derive coral cover of each category (Figure 3). Platform users were primarily volunteers, including the public, school children, and corporate staff partners in Corporate Social Responsibility programs. Users labelled polygons that were generated by the segmentation model described earlier. The label options were the same as for the expert analysis (Figure 2). A 3-minute video was provided when users first logged in to the platform to explain how to identify each category, with a help page available at all times. Floating pop-ups on the platform were also available on the image analysis page to remind users how to identify each group if required. For each analysis, the cover of each category was calculated in the same method as the expert and AI-only analysis; i.e. coral cover as a percentage of colonisable area in the image. When multiple users analysed the same image, the average of all user results was used.

**Figure 3.**
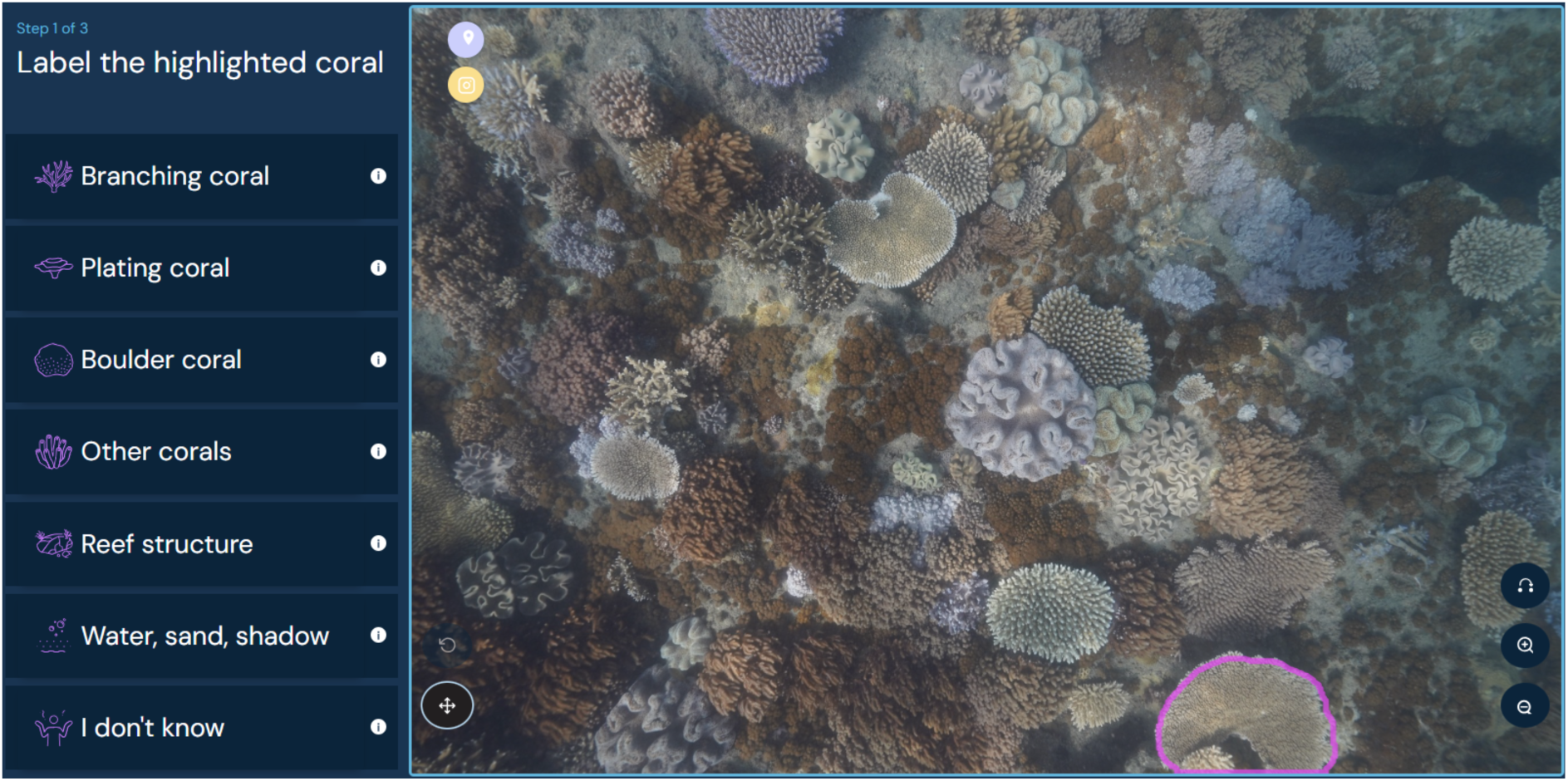
The Great Reef Census online analysis platform. Citizen scientists assigned labels to polygons generated by a segmentation model identifying distinct objects. The highlighted polygon to label can be seen in the bottom right corner. Credit: greatreefcensus.org.

The platform randomly assigned images to users in real-time, prioritising images with the fewest analyses complete. For example, if some images had already been analysed by two other users, the platform would only present images to users that had been analysed once. All images with the lowest number of complete analyses were equally likely to be assigned to a user, so that the images from a site were analysed by several online users.

The online platform was operational for 11 months (April 2023 – March 2024), during which 150,391 analyses of 20,879 images – each analysed multiple times – were completed by 6,052 individual citizen scientists from 70 countries.

### 1.2. Data analysis

We conducted a series of tests to examine the effectiveness and reliability of the citizen science method for collecting coral cover data. Based on the accuracies of other common tools (Leujak & Ormond, 2007), we chose ±5% absolute difference from expert values as an ecologically relevant accuracy target for broadscale reconnaissance; for example to be useful for distinguishing healthy from unhealthy reefs. To combine the relative strengths of the AI-alone and the AI+Citizens methods, the most accurate analysis method for each coral type was used in a ‘best’ method for all images. Next, while the mean of all images might be accurate, some management applications require coral cover estimates specifically at unhealthy reefs, in which case the method needs to be tested for images with low coral cover (0-20% coral cover). We disaggregated images into coral cover bins of 10% increments for each coral type, so that results can be interpreted within a diversity of reef contexts. For example, citizen science may overestimate low coral cover images and underestimate at high coral cover: a common problem for bounded proportion (0 – 100) metrics (Ferrari & Cribari-Neto, 2004). Consequently, any such bias may systematically over- or underestimate coral cover at individual locations. We then used simulations to determine how many images are needed to ensure a site estimate reliably falls within ±5% accuracy. This is required because although the mean value of all images may be accurate, there is variability in the accuracy of coral cover derived from any one image. Greater variability in accuracy among images will require more images from each site to obtain a reliably accurate mean site value. Finally, we performed power analyses to determine how many images are needed from a site to detect a 10% difference in coral cover, with 80% power, of each coral category.

#### 1.2.1. Accuracy of coral categories per image

##### 1.2.1.1. AI-alone

To determine the accuracy of the AI-alone method for each coral type, the mean expert result of each coral cover category *j* for each image *i* (*expert_ij_*, % cover) was subtracted from the AI-alone result of the same image (*AI_ij_*, % cover) to obtain an absolute percent difference 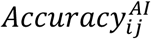(% cover):

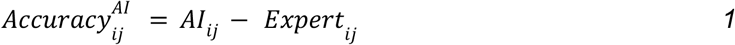

This was repeated for all coral categories. For example, if the AI output for Branching coral was 5% and the expert value was 10% for the same image, the AI-alone accuracy was described as-5%, i.e. AI underestimated the expert value by 5%.

##### 1.2.1.2. AI+Citizen

Similarly, to determine the accuracy of the AI+Citizen analysis for each image and coral category 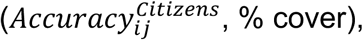 the mean expert result was subtracted from the mean AI+Citizen result to obtain an absolute percent difference:

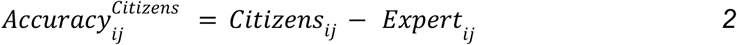

##### 1.2.1.3. ‘Best method’ accuracy

Given the relative strengths of the AI-alone and AI+Citizen results individually, we combined the results to achieve the ‘best’ method for analysing citizen science images. The best method used the more accurate-using the mean of all images-of the AI-alone or AI+Citizen method for each coral type 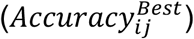 and applied it to all images.

##### 1.2.1.4. Disaggregating accuracy by reef state (coral cover)

To assess differences in accuracy at different coral cover levels, we categorised images into 10% cover bins for each coral category as determined by the experts. For each 10% bin with at least 80 images, we obtained the mean accuracy of images using our ‘best’ method for each coral cover category. Images were re-assigned to bins for each coral cover category.

#### 1.2.2. Images required per site

##### 1.2.2.1. Accuracy: Number of images needed to reach ± 5% accuracy

The earlier analyses provide the overall accuracy of the methodology in extracting coral cover from an image. However, given the variation in accuracy among images, we need to know how many images are needed for the mean accuracy of a site to meet an accuracy of ±5%. To answer this question, we ran a series of simulations.

For each simulation run, we randomly sampled *n* images from the entire image library and determined the mean accuracy 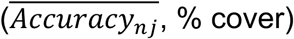 of each coral type *j* in those images:

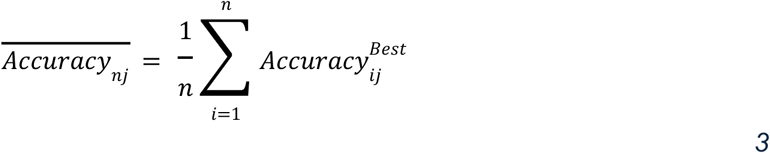

We conducted 10,000 simulation runs for each value of *n* from 1 to 120 and plotted each run’s value for 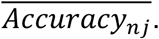.

##### 1.2.2.2. Effect of multiple citizen analyses per image

An advantage of the AI+Citizen analysis over AI-alone is that multiple citizen scientists can analyse the same image to obtain a mean result. The mean result from many individual analyses may be more accurate than having one citizen scientist analyse each image. As a result, if images are analysed by multiple citizen scientists, we may need fewer images to meet an accuracy of ±5% reliably, which can reduce the in-water survey effort. We assumed ‘reliably’ meant that an accuracy of ±5% is achieved in 95% of simulation runs. For the coral types for which AI+Citizen analysis was the most accurate, we determined the effect of increasing the number of analyses on the probability of a site being within ±5% of expert analysis. To achieve this, we repeated the simulations described in section 1.2.2.1 while varying the number of analyses per image (*m*) from 1 to 6. Analyses were sampled with replacement from each image. To obtain the mean accuracy of an image *i* with varying citizen scientist analyses (*v*):

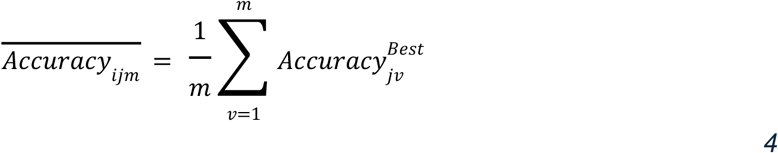

Where 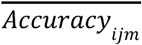 is the mean accuracy of coral category *j* for an image *i* with *m* number of citizen scientist analyses (*v*). We determined the mean accuracy across *n* images, given *m* citizen scientist analyses per image, by:

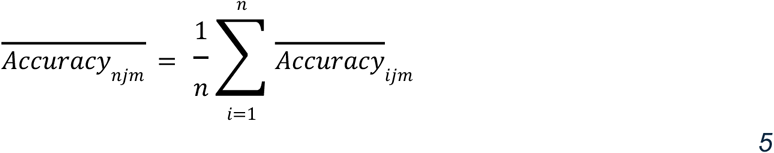

For each image count (*n* = 1 to 120) and analysis count (*m* = 1 to 6), the percent of runs that had a mean accuracy within ±5% was noted (out of the 10,000 runs for each combination of image count and analysis count). This provided the minimum number of images needed per site to meet an accuracy of ±5% in 95% of runs, to test if the number of images needed is reduced with more analyses completed per image.

##### 1.2.2.3. Power analysis: Number of images needed to detect 10% difference in coral cover

Once the minimum number of images to meet accuracy requirements for the methodology has been determined, there remains the question of capturing heterogeneity of the reefscape. A series of power analyses were performed to determine how many images per site are needed to distinguish between sites with a 10% difference in coral cover.

Images analysed by all methods (AI-alone, AI+Citizen and experts) were grouped according to survey site. Each site was categorised into 10% coral cover bins (0- 10%, 10-20% etc) for each coral type according to expert values. The standard deviation of coral cover values at each site was determined for each coral type using our ‘best’ method to capture the variability when using citizen science methodology. Then, within each coral type and coral cover bin, the mean standard deviation of coral cover at all sites was calculated.

The mean standard deviation of sites for each coral type and coral cover bin was used to conduct a power analysis, aimed at determining the minimum number of images needed per site to detect a 10% absolute difference in coral cover (effect size) with a power of 0.8 and an alpha level of 0.05. Any sites with fewer than 10 images analysed were discarded for this analysis. Images were not mixed across sites to ensure the realistic heterogeneity of the reefscape was captured.

All statistical analysis was performed in R (R Core Team, 2010) (R Core Team 2023) and the *tidyverse* collection of packages (Wickham et al., 2019). The power analyses were performed using the *pwr* package (Champely, 2020).

## 2. Results

### 2.1. Accuracy of Coral Categories per Image

#### 2.1.1. Mean accuracy of AI-alone and AI+Citizens

The mean difference between the expert analysis and AI-alone analysis for all images (8,086 images) ranged from-9.1% for Plating coral to +6.9% for Other coral (Figure 4). The mean difference between expert analysis and AI+Citizen analysis for all images with at least 1 citizen analysis (7,790 images) ranged from-0.99% for Plating coral to +9.5% for Branching coral (Figure 4).

**Figure 4.**
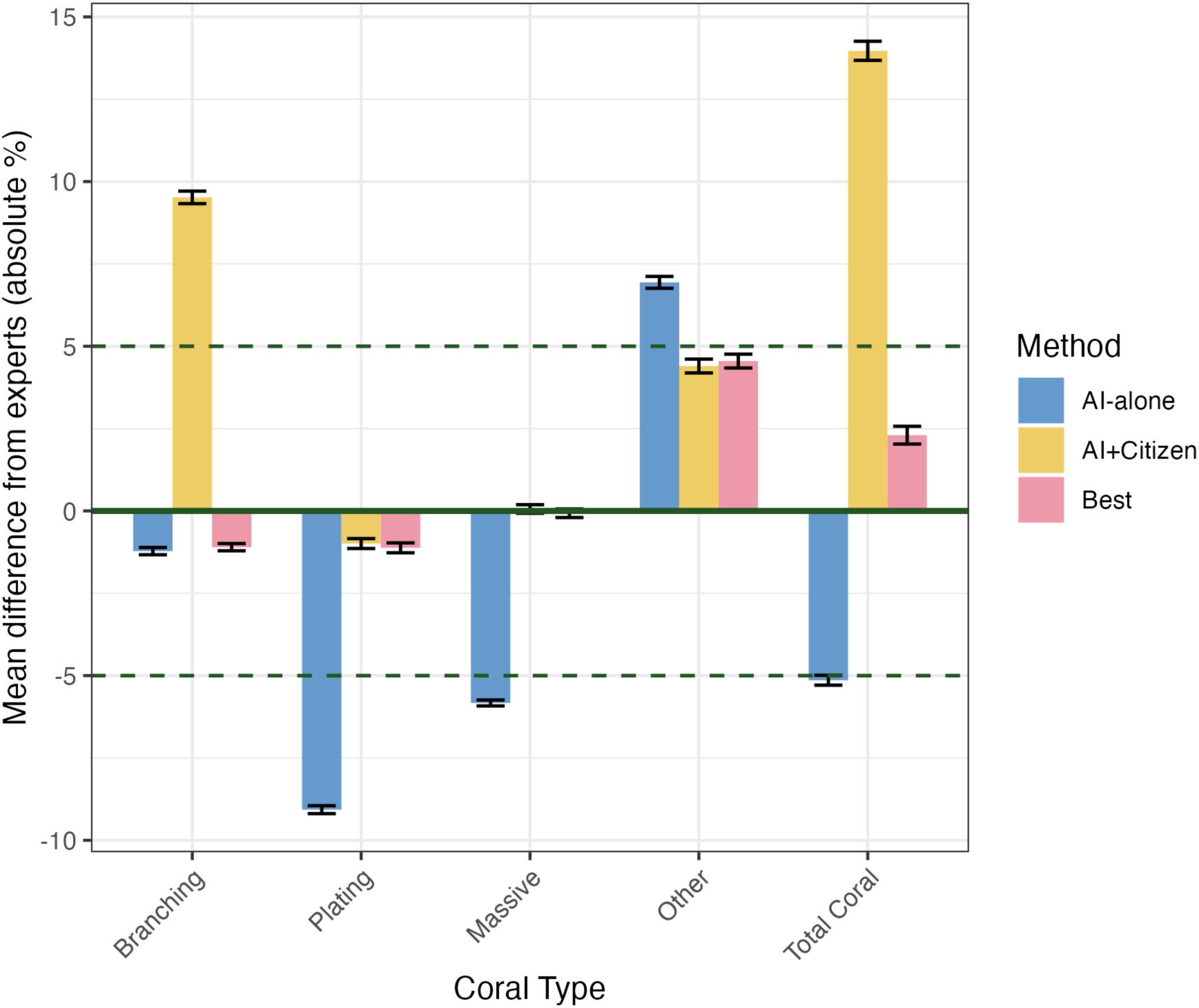
The mean accuracy of the AI-alone (7,505 images), AI+Citizen (7,790 images) and ‘Best’ (7,608 images) method for each coral category. The y-axis is measured as the difference between the method’s output and the expert results for each same image. “Total Coral” is the accuracy of the total benthic coral cover, i.e. the sum of the difference from expert analysis of all individual coral categories. “Other” refers to all coral types except branching Acropora, plating Acropora, and massive-form corals. Error bars show standard error of the mean. NB: Negligible differences are observed between the best method and the most accurate method for each coral type (e.g. AI-alone and best for Branching coral) due to slight differences in which images were analysed for each method.

The AI-alone method was more accurate for Branching coral cover, while the AI+Citizen method performed better for Plating, Massive, and Other coral cover. Therefore, the ‘best’ method combined AI-alone results for Branching coral with AI+Citizen results for the remaining coral types. The mean difference from experts using our best method was-1.1% for Branching coral, -1.1% for Plating coral, -0.1% for Massive coral and +4.55% for Other coral. Using our ‘best’ method, the mean difference from experts for total coral cover improved from -5.1% (AI-alone) and +13.9% (AI+Citizen) to +2.3% (Figure 4).

#### 2.1.2. Disaggregating accuracy by reef state (coral cover)

For Branching, Plating and Massive coral, all reef state bins were within our target of ±5% accuracy (Figure 5), but Other coral had higher error for low and high reef state bins, ranging from +9.3% for 0-10% coral cover to -19.7% for 40-50% coral cover (Figure 5). There was higher uncertainty in mean accuracy at high coral covers due to small sample sizes (Figure 5).

**Figure 5.**
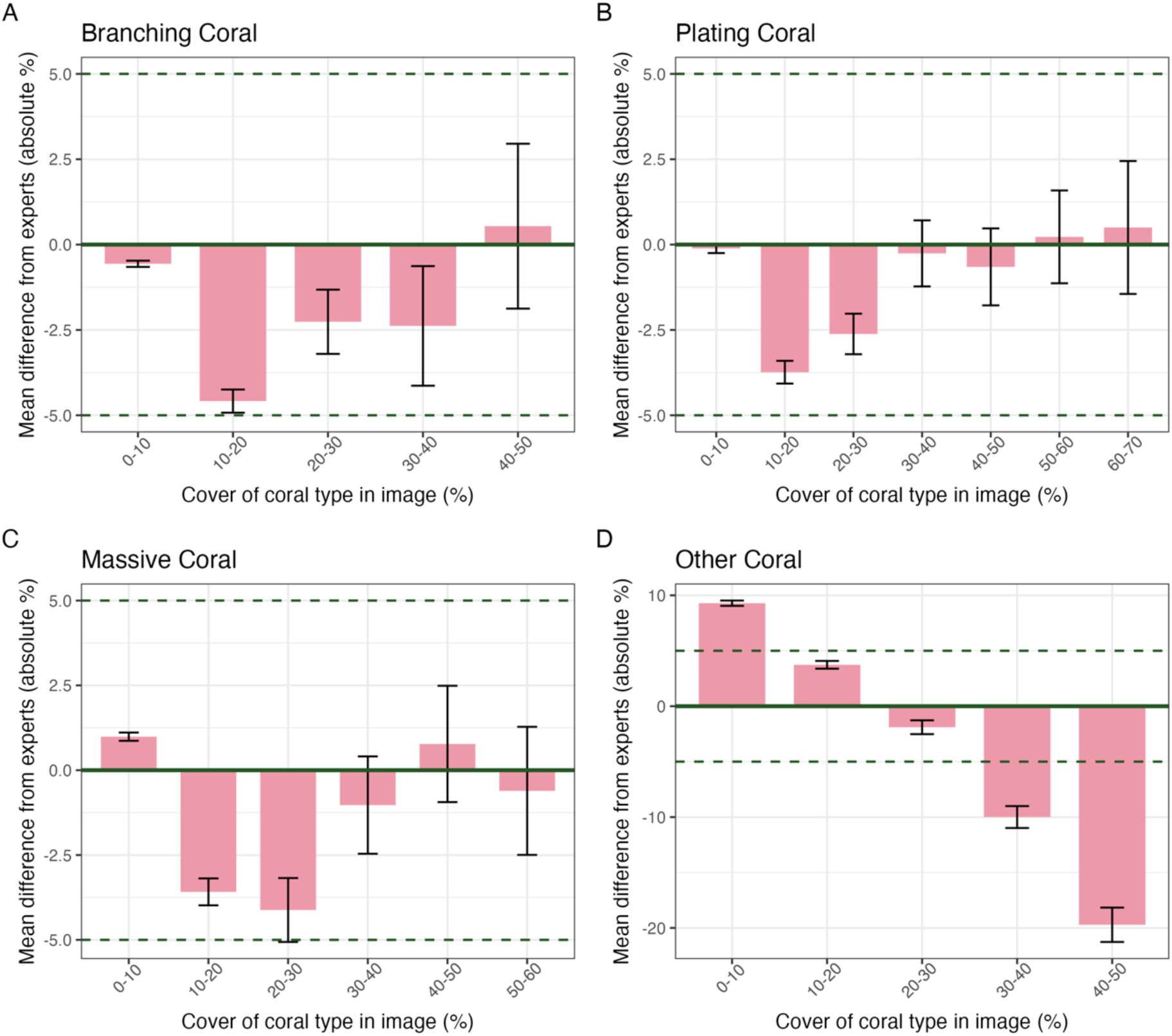
The mean accuracy of coral cover estimates for images from each 10% reef state bin using the ‘best’ method (minimum images per reef state bin = 80, because up to 80 images were needed to achieve accuracy in 95% of sites for all coral types with one citizen analysis complete; see Figure 6 Plating coral panel). The x-axis represents the coral cover of the coral category according to expert analysis. Error bars show standard error of the mean; there were generally fewer images available at higher coral cover bins, resulting in larger standard errors. Dashed horizontal lines show our desired accuracy threshold (± 5%). Note that the y-axis range differs in panel D.

### 2.2. Images required per site

#### 2.2.1. Accuracy: number of images needed to reach ± 5% accuracy

The simulations showed that increasing the number of images per site reduced the variability in mean site accuracy (Figure 6). For example, with just one image per site, 95% of sites had differences from expert analysis ranging from -11% to +22% in absolute Branching coral cover. In contrast, when 80 images were collected per site, 95% of sites showed differences within a narrower range of -3% to +1%.

**Figure 6.**
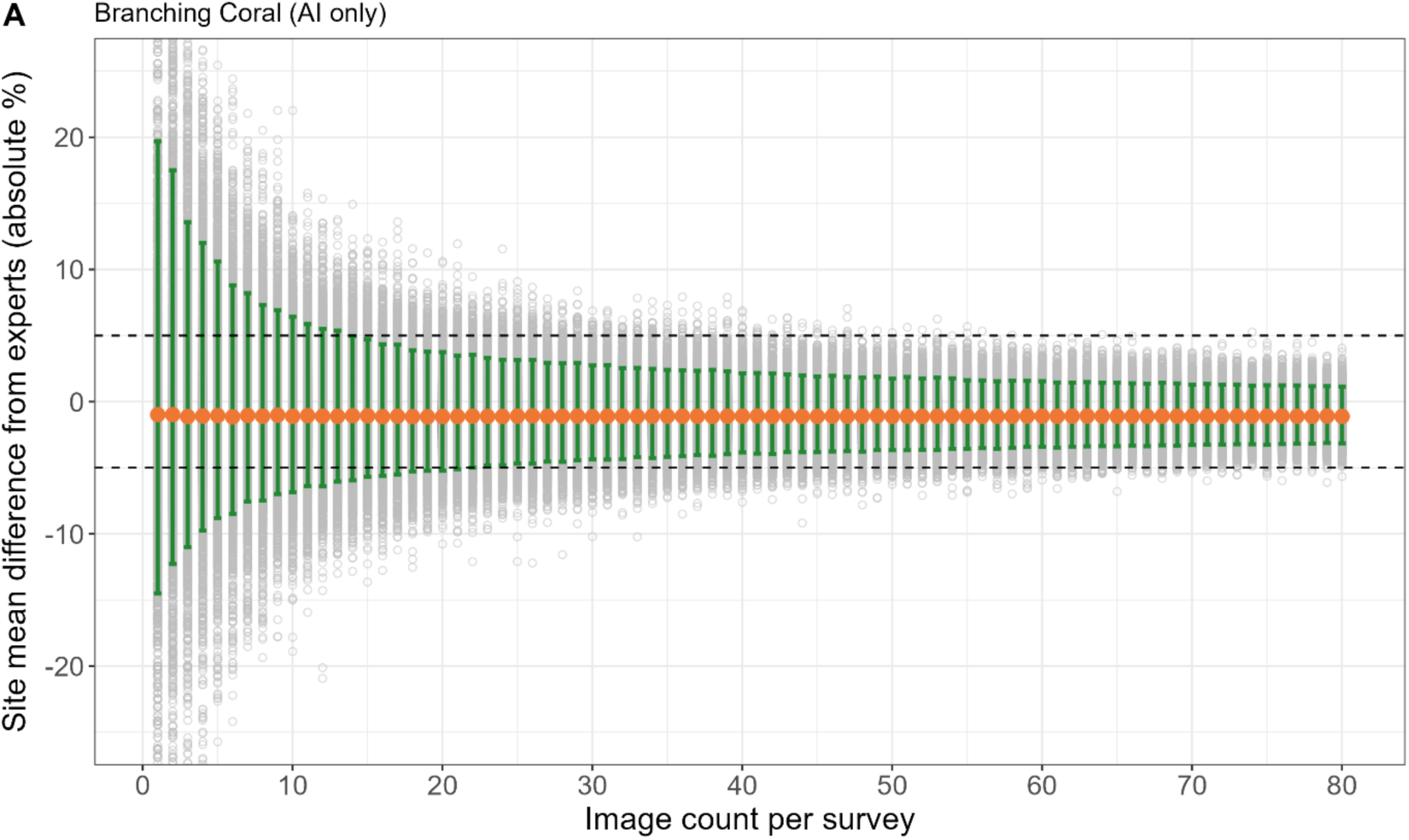
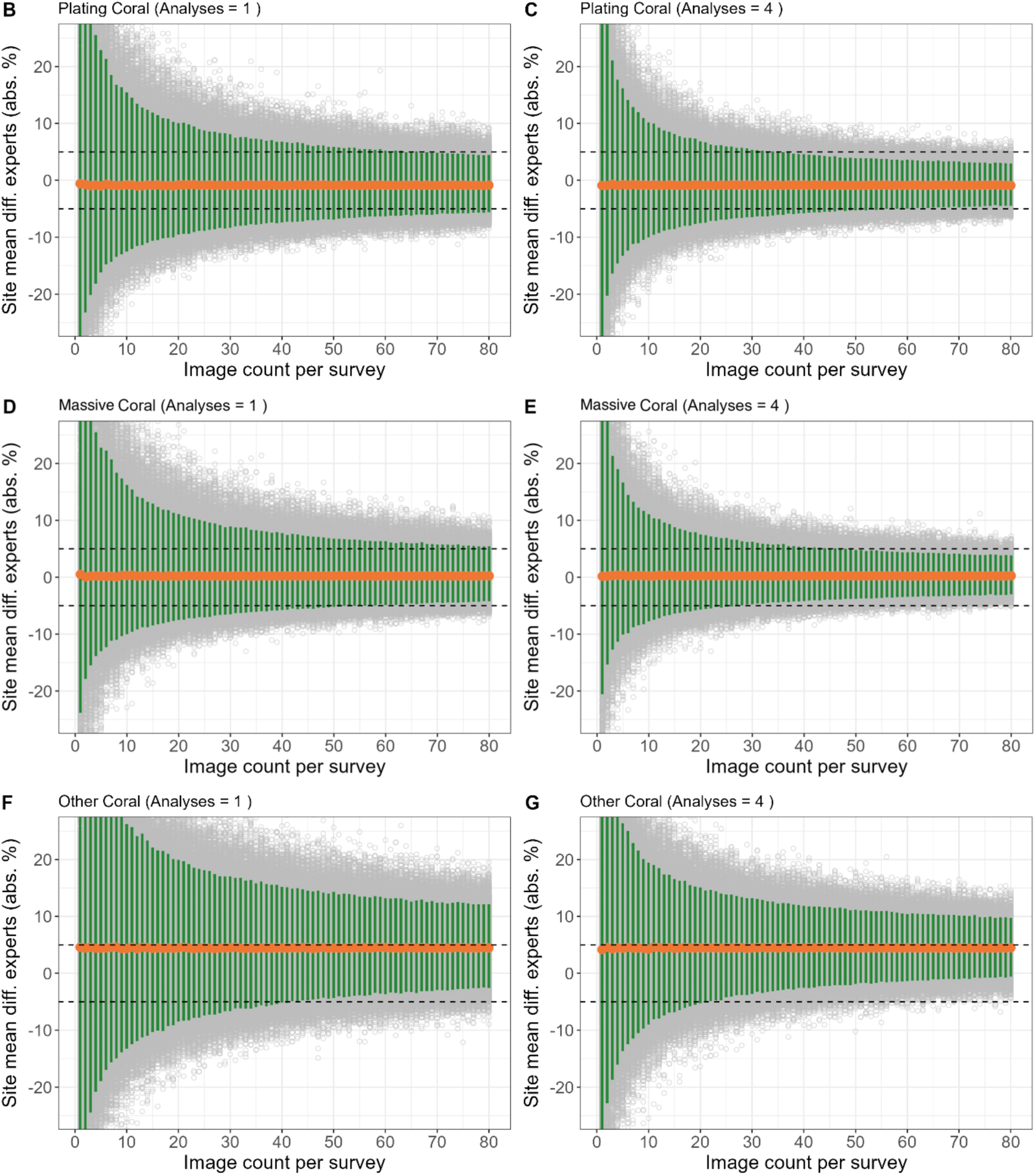
Mean site accuracy with increasing image count for Branching coral using AI-alone (A), Plating coral with 1 citizen analysis per image (B), Plating coral with 4 citizen analyses per image (C), Massive coral with 1 citizen analysis per image (D), Massive coral with 4 citizen analysis per image (E), Other coral with 1 citizen analysis per image (F) and Other coral with 4 citizen analysis per image (G). Each grey point represents the mean image accuracy of one simulation run of randomly sampled images (10,000 runs per image count value). The orange points represent the mean value of all simulation run means for each image count value. The green bars show where 95% of simulation runs lie. The simulations were run up to 120 images per survey site, but the x-axis is truncated for clarity here.

Consequently, collecting more images from each site increased the likelihood of the site accuracy meeting an accuracy of ±5%. Branching coral – the only category in which AI-alone was most accurate – required 17 images per site for the mean accuracy to be within ±5% accuracy for 95% of sites (Figure 6a; Figure S1). Plating and Massive coral needed less than 80 and 70 images, respectively, to achieve ±5% accuracy for 95% of sites, but this varied depending on the number of citizen analyses completed on each image (see later). For Other coral, as the number of images collected per site increased, the percent of sites that achieved ±5% accuracy became asymptotic to about 60% (Figure 6).

#### 2.2.2. Effect of multiple citizen analyses per image

For coral categories in which AI+Citizen was more accurate than AI-alone (Plating, Massive and Other), increasing the number of analyses per image reduced the number of images needed per site to achieve ±5% accuracy, with diminishing returns (Figure 6b-g). For example, with just one analysis per image, 80 (Plating) and 70 (Massive) images were needed to meet an accuracy of ±5% for 95% of simulated sites, yet if 4 analyses were completed then just 44 and 34 images, respectively, were needed (Figure 6b-g; Figure S1). Completing 6 analyses per image only marginally reduced the required images to 40 and 31 images for Plating and Massive categories, respectively. In general, 4 analyses per image achieved high accuracy with efficient resource use, however this will vary depending on project goals and resource distribution across in-water survey and online analysis efforts.

#### 2.2.3. Power analysis: number of images needed to detect 10% difference in coral cover

The power analyses showed that the number of images required to detect a 10% difference in absolute coral cover ranged from 4 (Branching coral 0-10%) to 114 per site (Massive coral 30-40%; Figure 7). Most of the tested categories required 80 images or less to detect a 10% difference in absolute coral cover of that category. Generally, more images were needed at higher coral covers. Few sites were available with coral cover greater than 50% in any coral category.

**Figure 7.**
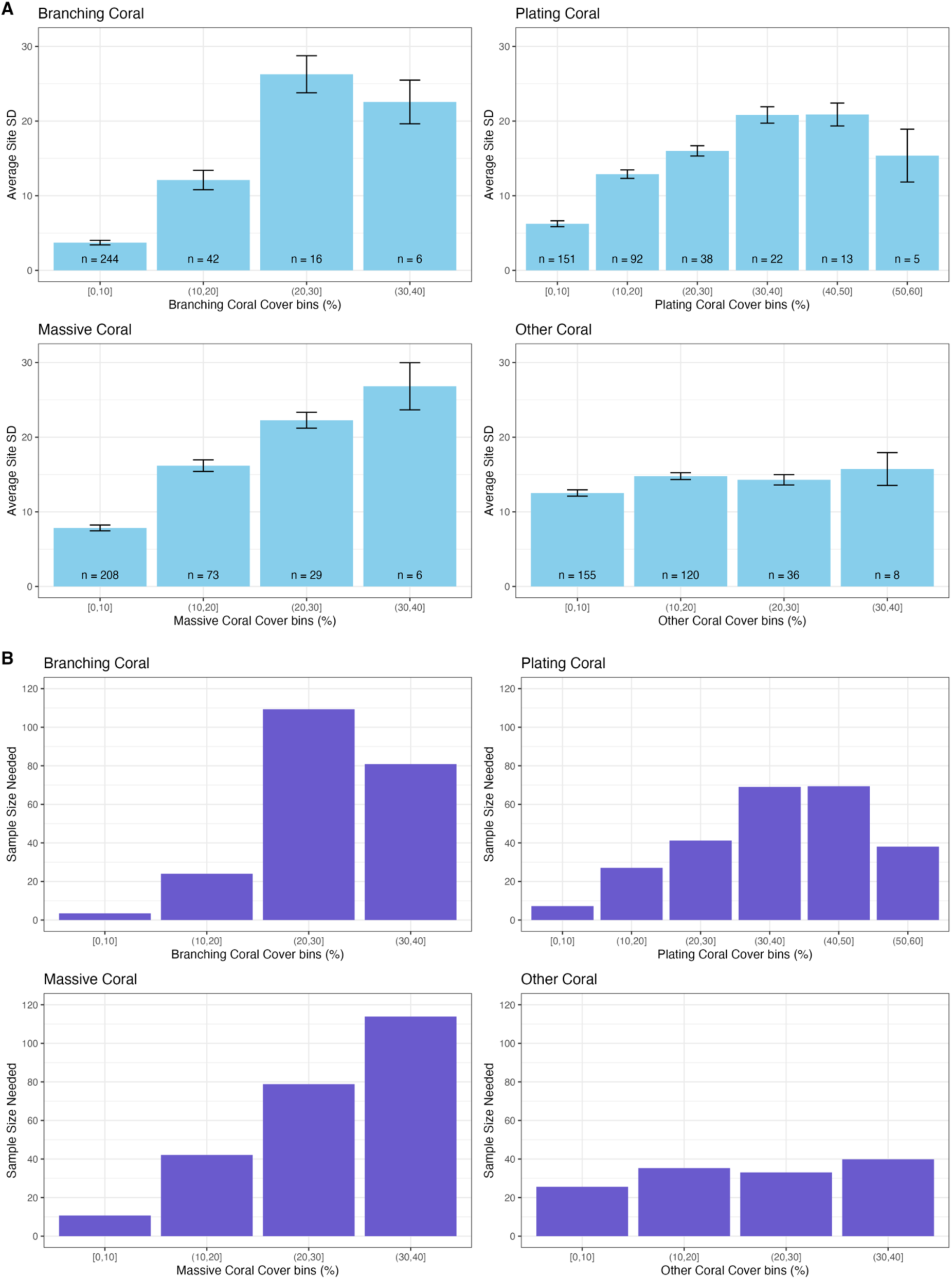
A: Mean standard deviation of surveyed sites for each reef state bin and coral type. Error bars show standard error of the standard deviation. n-values inset show the number of sites in each column. B: Power analysis results. Columns show the number of images required per site to detect a 10% difference in coral cover among sites based on their standard deviation (power = 0.8, alpha = 0.05).

## 3. Discussion

A combination of AI and non-expert human analysis of seascape images collected by citizen scientists can provide cover estimates of key coral categories that are accurate to within ±5% of trained expert analysis. This accuracy was achieved at any level of coral cover for Branching, Plating and Massive coral, but was only achieved for Other coral in images with 10-30% cover. The level of citizen science effort required to meet ±5% accuracy for the three key coral categories – up to 45 images per site analysed by four citizen scientists – is achievable based on previous participation in citizen science initiatives. Power analyses demonstrated that for some sites, more images are needed to detect a 10% change in coral cover and capture the heterogeneity of the reef than are necessary to be confident in the accuracy of the analysis method. Here we discuss the practical application of these methods and considerations dependent on project goals.

### 3.1. Varying the sampling protocol based on project goals

A project using a citizen science-based method similar to that presented here can adjust its sampling strategy based on the program goals and distribution of resources between in-water survey efforts and online citizen scientists (Table 2). If more resources are allocated to online citizen scientists than in-water sampling, the project could reduce the number of images collected, relying on increased citizen scientist analysis effort to maintain confidence in the results. Over the first two years of testing the online analysis platform, each image was analysed 5-6 times. The platform’s scalability suggests that this level of analysis can be sustained given that online analysis is cheaper and can be conducted globally, while in-water surveys require more resources and are restricted to local participants. Indeed, in some instances collecting fewer images per site and surveying more sites is a preferred approach, as more extensive online citizen science analysis could compensate for the lower image count.

**Table 2.** Example scenarios to illustrate the interplay between the number of images required to meet methodological accuracy levels and the number of images required to detect a 10% difference in coral cover between sites based on the results of the power analysis. Fewer images will decrease the certainty in coral cover estimates, however this may be acceptable for some project goals. In most cases, the required number of images to meet desired methodological accuracy will need to be met, regardless of if the results of the power analysis need to be met. However, in some instances, a ± 5% accuracy target is higher than required and so fewer images can be collected to meet a lower accuracy target.

| Scenario | Required images relative to number needed for $\pm 5\%$ methodological accuracy. | Required images relative to number needed to detect a 10% difference. |
| --- | --- | --- |
| Low precision needed: 15-20% detectable difference in coral cover between sites. | Similar | Fewer |
| High precision needed: 5% detectable difference in coral cover between sites. | Similar | More |
| Approximate coral cover needed to distinguish very low and high coral cover reefs. | Fewer | Fewer |
| Outplant restoration site with low, homogenous coral cover. | Similar | Less than needed to meet $\pm 5\%$ accuracy. |
| Validate habitat maps of dominant coral type. | Similar | Likely fewer. |

For example, there is management interest in validating modelled habitat maps of key coral morphologies (Roelfsema et al., 2021). These maps predict the coral morphology most likely to dominate based on environmental factors such as wave energy and disturbance exposure. Such maps support research, ecological modelling, and decision-making in management and restoration (Anthony et al., 2017; Bellwood et al., 2019; Pittman et al., 2007). However, the modelled predictions of dominant coral type often lack empirical validation. To validate these maps most effectively, it is essential to survey as many sites as possible given that dominant coral type can vary over short distances. Hence, using online citizen science analysis to improve accuracy and minimise image collection at any one site is preferred.

Sampling design can also be guided if the approximate condition of the reef is known *a priori*. For example, if a site is known to be heavily damaged with less than 20% coral cover, then the power to detect change is unlikely to be an issue if enough images are collected to meet accuracy needs for the sampling method (generally at least ∼44 images with 4 analyses each). A similar approach may be taken to survey small scale restoration activity where most of the area can be surveyed directly and/or is likely to be highly homogenous (McLeod et al., 2022).

Similarly, if a project needs less accurate estimates of coral cover, say within ±10%, fewer images are needed to be confident in the method. As coral cover increases, it is likely less important to obtain a highly accurate and precise estimate of coral cover. For example, a ±10% range in possible values at 50% coral cover is unlikely to affect decision-making in the same way it would at 15% coral cover (Wickham et al., 2019), unless the goal is to track coral cover change precisely over time.

### 3.2. Samples required compared to other tools

Here we showed that useful broadscale reconnaissance survey data can be achieved with currently observed levels of citizen scientist engagement. In some situations, such as sites with Branching 20-30% and Massive coral 30-40%, more images were needed to detect a 10% difference in coral cover with sufficient power than were needed to be confident in the accuracy of the sampling method. The limiting factor at such sites may be the natural heterogeneity of the reef rather than the accuracy of the sampling method. This is reflected in traditional reef surveying methods such as photo quadrats and line transect point methods, which require sampling similar to or greater than needed here. For example, to detect a 20% relative difference in coral cover using photo quadrat methods, above 10% absolute cover, requires 38 - 48 (branching *Acropora*) and 111 – 141 images (massive *Porites*), or using line transect methods requires 990 - 15,450 (branching *Acropora*) and 820 – 8200 points (massive *Porites*) (Leujak & Ormond, 2007). Similarly, Carneiro et al. (2024) found that substantially more survey effort was required to achieve equivalent accuracy and precision by two common line transect survey methods, Reef Check and the Atlantic and Gulf Rapid Reef Assessment, compared to photo quadrats. To estimate coral cover with a 20% error margin, Reef Check required 1280-3080 line transect points and Atlantic and Gulf Rapid Reef Assessment required 1400-2200 line transect points (Carneiro et al., 2024).

The distribution of effort among the number of images collected per site, sites surveyed, and analyses completed per image will depend on the resource availability and goals of a program. However, the approximate requirements presented here are achievable based on experience. For example, while collecting 80 images per site (40 images each by two snorkellers), previous Great Reef Census expeditions with four participants have surveyed up to 124 sites across 42 reefs in six days (pers. comm. A. Ridley, Citizens of the Reef). Similarly, in the first two years of the Great Reef Census operating, all images (up to 29,967 per year) have been analysed by at least 5 online citizen scientists with participants from 80 countries (unpublished data). Given this observed effort and the potential for widespread use by citizen scientists, such a method may expand data collection in resource-poor areas or provide an efficient complement to existing methods (Madin et al., 2019).

### 3.3. Correcting for known inaccuracy

If there are systematic biases that cause known inaccuracies in a method, a correction offset can be included when reporting results (e.g. Eikelboom et al., 2019). For example, a 5% methodological overestimation may reduce the data’s reliability for management decision-making. Hence, any estimates of accuracy can be used as an offset to correct the data.

Here, applying a constant offset is likely suitable for Branching, Plating and Massive coral estimates because all coral cover bins for these categories had similar accuracies that were reliably within ±5% of the expert analysis. Applying such an offset should not affect the uncertainty of estimates, and therefore will not affect required sample size, because the offset is an absolute percentage of a proportion rather than a relative percentage offset (Eikelboom et al., 2019). However, care should be taken if applying an offset for Other coral results, which had more variable accuracy depending on coral cover level. Other coral was overestimated at low coral covers and underestimated at high coral covers, making it difficult to apply a constant offset. This may be a limitation of the current method, in that accurate estimates of cover can be provided for Branching, Plating and Massive coral but total coral cover will be underestimated at sites with high Other coral cover.

### 3.4. Future improvements and conclusions

The main drivers of improved performance in distributed data collection and analysis programs will likely be technological, although improved training of citizen scientists and program design can help. For example, anecdotally, we observed that poor quality images appeared to be harder for both the AI and citizen scientists to analyse accurately. Poor quality images were commonly caused by human/camera error, poor water visibility, or images captured more than 5 m from the reef. As camera technology improves and becomes cheaper, the occurrence of poor-quality images will likely reduce. Similarly, participants could be instructed to capture images closer to the sea floor, for example at 3 m instead of 5 m, especially in poor water visibility. Improved access to post-processing tools, such as automatic colour correction, can also improve image quality (Raveendran et al., 2021). These factors, alongside improvements in segmentation model technology, will make analysis by AI and humans easier and likely improve accuracy. In terms of training citizen scientists, clearer instruction for identifying dead coral may improve accuracy. Dead Branching coral in particular – the only coral category for which AI-alone was more accurate than AI+Citizens – appeared to be poorly identified (pers. comm. Citizens of the Reef).

Major improvements may also be achieved by increasing the number of benthic categories that can be accurately measured. The Other coral group here was the least accurate likely because it encompasses all coral types except our three key morphologies, making segmentation model training difficult (Rubbens et al., 2023). The uncertainty in Other coral estimates may be resolved by disaggregating the category into distinct coral morphologies/taxonomies and through continual advances in deep learning (González-Rivero et al., 2020). More resource-intensive citizen science programs can assess dozens of benthic categories (Done et al., 2017) and emerging deep learning software can identify some coral to the species level (González-Rivero et al., 2020). However, there is a trade-off between data quality and scalability; higher taxonomic resolution data currently requires high quality photographs or participant training that intrinsically limits the program’s potential span of data collection.

A program such as the Great Reef Census demonstrates how technology, particularly deep learning, can lower the barrier to entry for citizen science, allowing non-experts to contribute to accurate coral reef data collection. This approach can enable large-scale participation globally. While not a replacement for more detailed scientific monitoring, the method may provide a complementary tool that can support coral reef management, especially in resource-limited regions, by offering an accessible and cost-effective method for broadscale surveying of key coral morphologies.

## Acknowledgements

This work was funded by the Australian Government under the National Environmental Science Program. We thank the thousands of citizen scientists globally who contributed to in-water surveying or online analysis, as well as Andy Ridley, Ben Vozzo, Jasmina Uusitalo and Nicole Senn from the not-for-profit organisation Citizens of the Reef who lead the Great Reef Census. We also thank the dozens of high schools and companies that donated student/staff hours to online analysis, including Dell Technologies, Disney Voluntears, Cotton On Group, Mars, Microsoft, Salesforce, Atlassian, PADI, and Virgin Voyages. Thank you to the Traditional Owner groups along the Great Barrier Reef who collaborated with us to work on their Sea Country, and the many tourism and marine industry operators who helped collect images. Thanks to our ‘experts’ for their diligent image analysis, Chandru Ganesan and the team at Sahaj Software Solutions for building the online platforms, and the team at Dell Technologies for developing the segmentation model as part of a Singapore Government research grant. The Great Reef Census is supported with donations and in-kind support from The Cotton On Foundation, The Prior Family Foundation, The Walt Disney Company, Mars, Mindshare, Cairns Regional Council, Reef and Rainforest Research Centre, The Australian and Queensland Governments, The Triumph Conservation Trust, and many individual donors.

## Notes

### Competing Interest Statement

The authors have declared no competing interest.

