## Supplementary material for "Broadscale reconnaissance of coral reefs from citizen science and deep learning": Figure S1

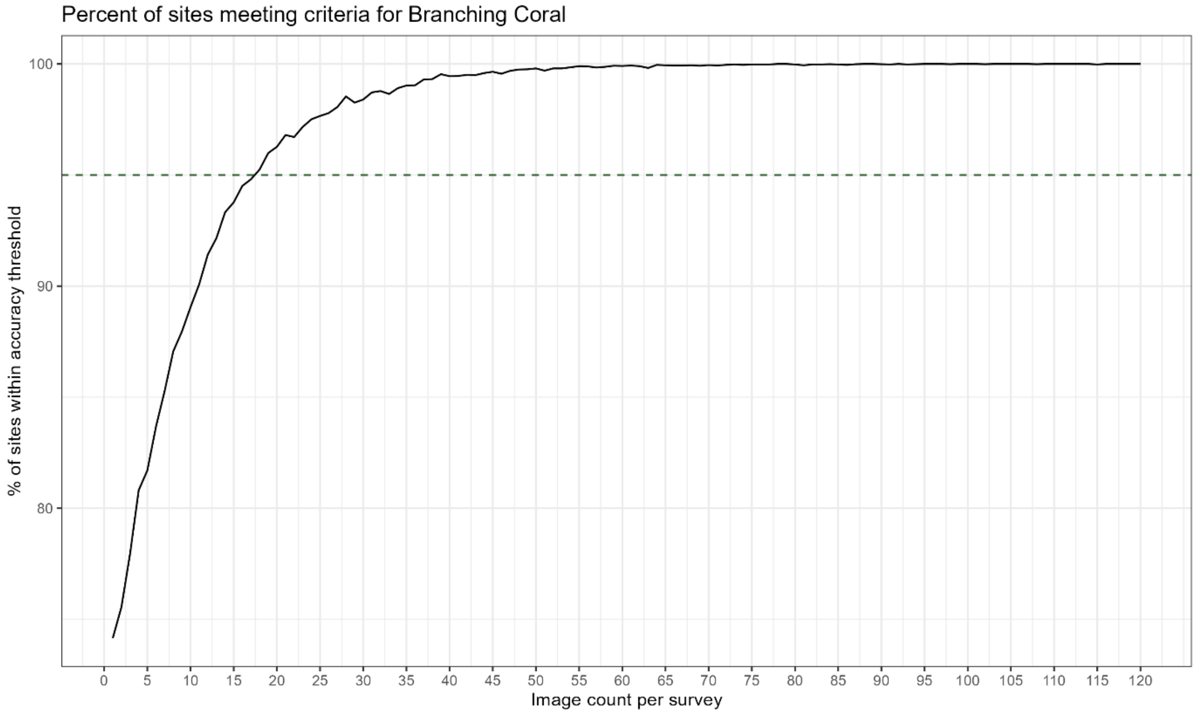

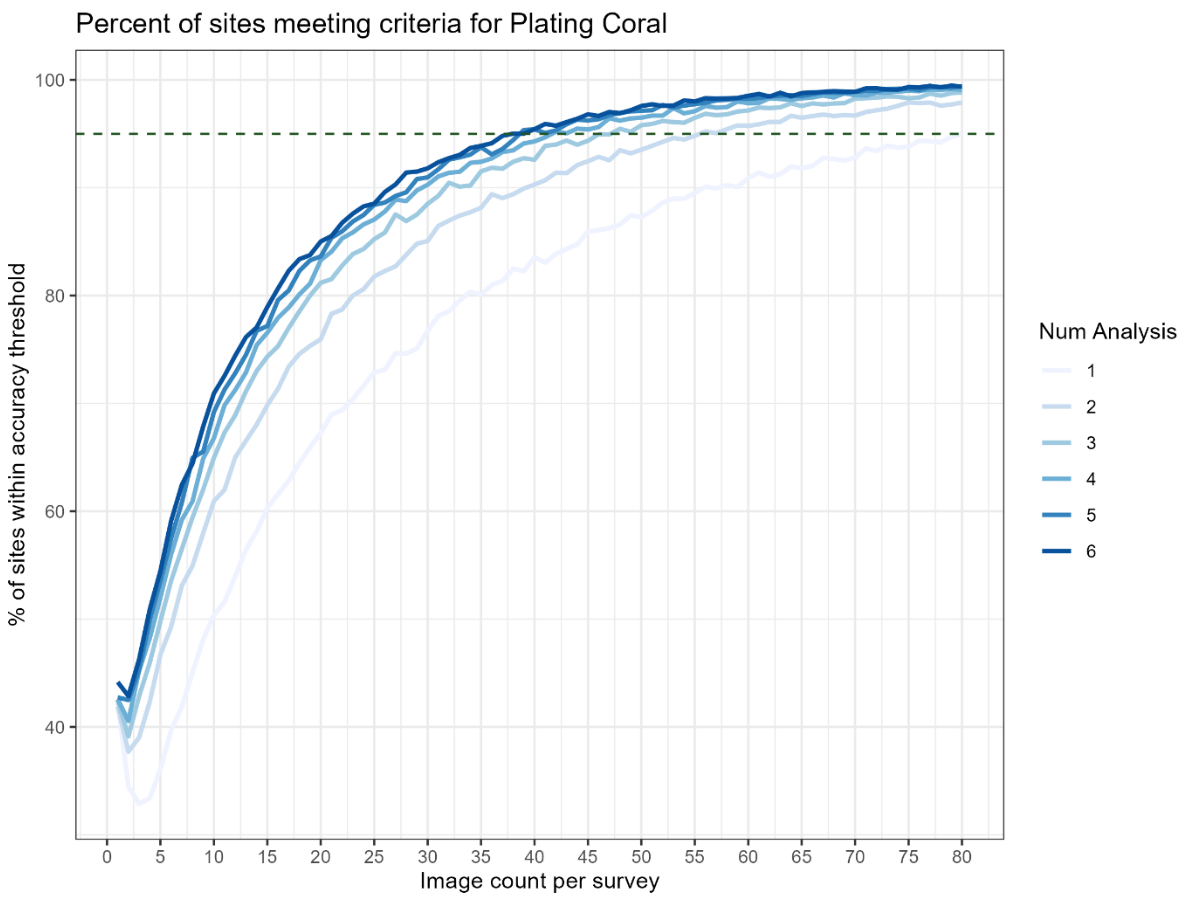
Percent of sites meeting criteria for Plating Coral
% of sites within accuracy threshold
100
80
60
40
0
5
10
15
20
25
30
35
40
45
50
55
60
65
70
75
80
Image count per survey
Num Analysis
1
2
3
4
5
6

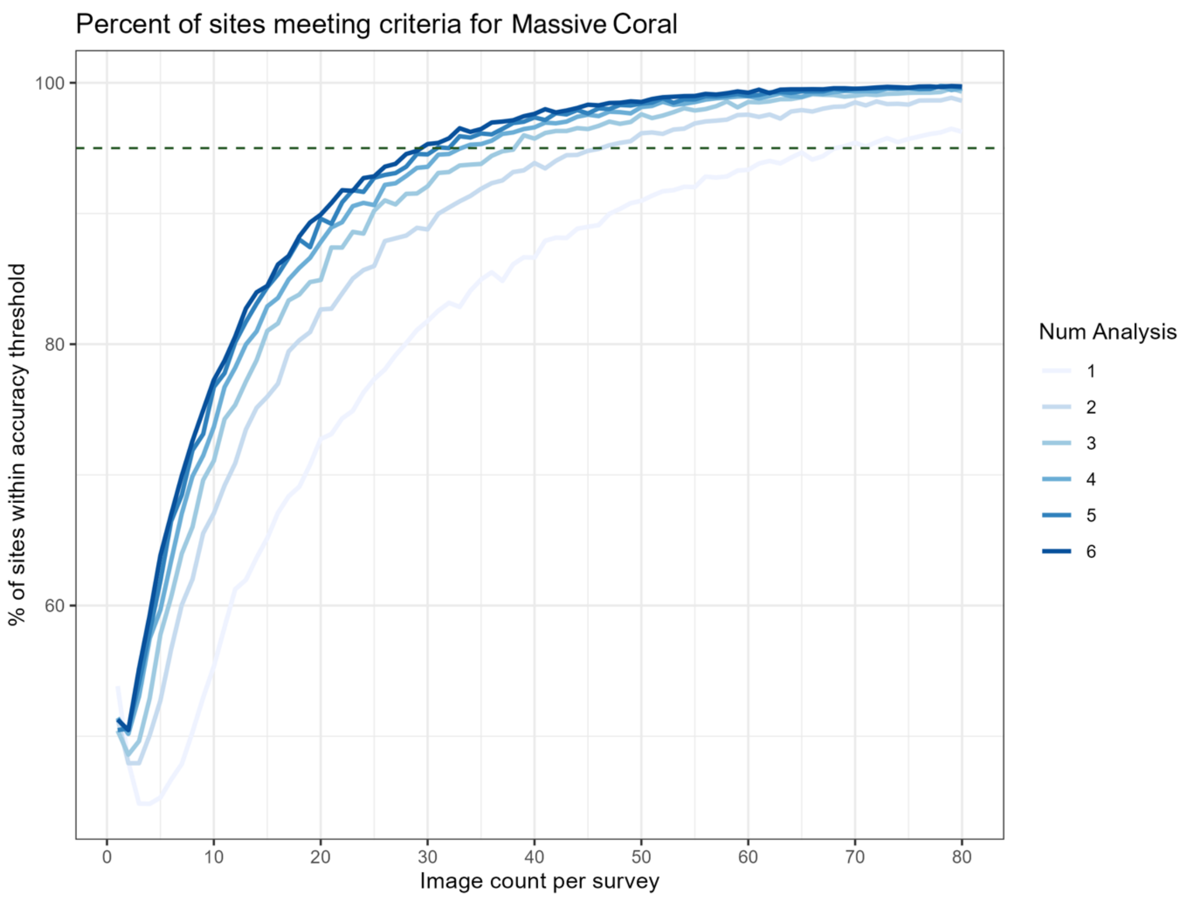
Percent of sites meeting criteria for Massive Coral
% of sites within accuracy threshold
100
80
60
0
10
20
30
40
50
60
70
80
Image count per survey
Num Analysis
1
2
3
4
5
6

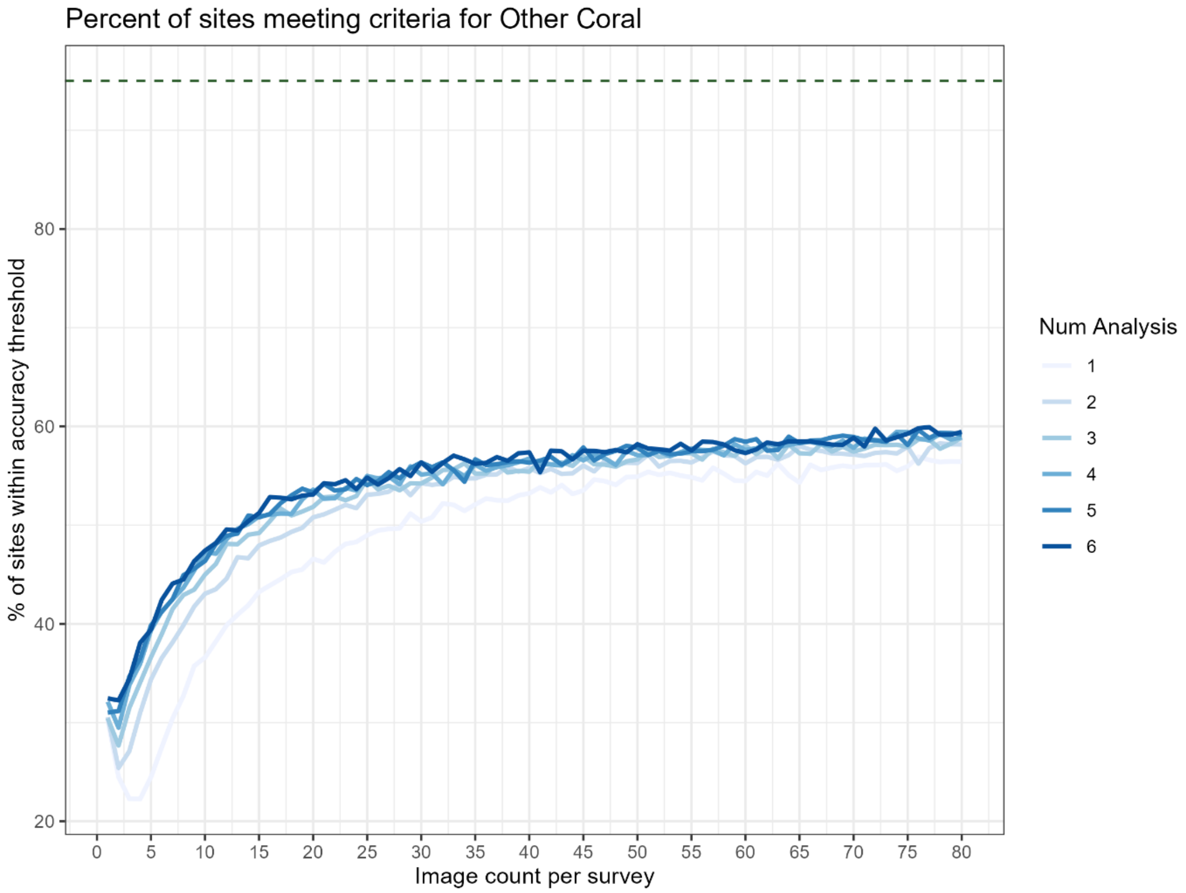


*Figure S1 – Cumulative percent of simulated sites that meet ±5% mean accuracy relative to expert analysis. Branching coral shows the results of AI-alone. The remaining categories show results from the AI+Citizen online analysis, with each line representing the results of varying the number of citizen analyses performed on each image. The dashed horizontal line shows 95% of simulated sites.*
